# High-level visual and motor regions are doubly dissociated in controlled tasks but only at specific times during natural behaviors

**DOI:** 10.64898/2026.09.29.755544

**Authors:** Shubhankar Saha, Sini Simon, Surbhi Munda, Thomas Cherian, Georgin Jacob, Jhilik Das, Debdyuti Bhadra, SP Arun

**Author notes:** indicates equal contributions (Sini Simon, Surbhi Munda, Thomas Cherian).

## Abstract

Our knowledge of the neural basis of sensory and motor information processing comes largely from recording brain activity in controlled settings. This approach, while insightful, blinds us from understanding natural behaviors, which involve far richer visual inputs and far larger range of movements. The prevailing belief is that, during natural behaviors, movement information would modulate visual cortex, and visual inputs would modulate motor cortex. Alternatively, this mixing might only reflect correlations between sensory inputs and movements present during natural behavior. To resolve this issue, we performed wireless recordings from high-level visual (IT) and motor regions (PMv/PFC) in freely moving monkeys while they performed a natural task and a fixation task. In each task, we extracted visual and movement features and asked to what degree these features explained neural activity. In the fixation task, we found a striking double dissociation whereby visual (but not movement) features predicted neural activity in IT, whereas motor (but not visual) features predicted neural activity in PMv & PFC. This dissociation was far weaker in the natural task but present particularly at key events when the visual and motor features were less correlated. Thus, high-level sensory and motor regions in the brain are doubly dissociated under controlled conditions, and remain so at specific times during natural behavior.

## INTRODUCTION

Natural behaviors have long been considered extremely challenging to study, since they involve large variations in both sensory and motor information. The simple act of eating food can involve making saccades, seeing the food from different angles each time, and reaching differently each time. Partly for this reason, and partly due to methodological limitations, traditional neuroscience paradigms have recorded brain activity under artificial conditions where many confounding variables are removed or minimized. While this approach has been extremely insightful (Rust and Movshon, 2005), it also prevents a deeper understanding of brain activity during natural behavior (Miller et al., 2022; Cisek and Green, 2024; Leopold, 2024; Parodi et al., 2025). With the advent of wireless recording technologies, brain activity in monkeys has now been recorded under increasingly natural conditions. In such conditions, prefrontal and motor regions in monkeys have been found to encode novel motor and navigational variables (Berger et al., 2020; Hansmeyer et al., 2023; Maisson et al., 2023; Voloh et al., 2023; Lanzarini et al., 2025), behavioral states such as grooming and sleep (Hsieh et al., 2008; Gabbott and Rolls, 2013; Yin et al., 2014; Walker et al., 2021; Bukhtiyarova et al., 2022; Yun et al., 2023) and even variables relevant for social interactions (Franch et al., 2024; Testard et al., 2024).

A fundamental unresolved question in this context is the extent to which sensory and motor signals are intermixed during natural behavior. The prevailing view is that since natural behaviors involve large changes in sensory inputs and motor movements, movement signals might strongly modulate sensory regions and conversely, sensory inputs might strongly modulate motor movements (Leopold, 2024; Kang et al., 2025). Alternatively, the mixing of sensory and motor signals during natural behavior might be simply due to sensory and motor signals themselves being correlated during natural behavior, in which case the mixing will disappear when these signals are uncorrelated. This view is supported by recent evidence that neural responses in primate visual cortex are relatively unaffected by small spontaneous movements in restrained animals (Talluri et al., 2023; Kang et al., 2025), although it is not clear how much these results will generalize to completely unrestrained naturalistic behavior. Addressing this issue will require recording brain activity under controlled conditions to establish that visual & motor cortex are indeed driven by visual and motor information, and then testing whether this dissociation holds during natural behavior.

### Overview of this study

Here, we set out to resolve this fundamental gap in our knowledge through wireless brain recordings in freely moving monkeys in controlled and natural conditions. We recorded simultaneously from the inferior temporal (IT) cortex, a high-level visual region that contains invariant visual object representations (Connor et al., 2007; DiCarlo et al., 2012; Conway, 2018) and from the premotor/prefrontal regions (ventral premotor cortex, PMv and ventrolateral prefrontal cortex, PFC), which contain invariant motor and task representations (Rizzolatti and Luppino, 2001; Schrijver et al., 2024).

In the natural setting, we recorded brain activity while monkeys housed in a large arena took food items offered by human experimenters. In the controlled setting, we recorded brain activity while monkeys performed a fixation task involving images of the same food and experimenters on a screen. This allowed us to evaluate the dissociation between visual and motor regions under controlled conditions and test whether this dissociation holds under natural settings.

## RESULTS

To understand the encoding of visual and motor information in controlled and natural settings, we tested each monkey in two sessions. In the touchscreen session (Figure 1A), the monkey had to perform a fixation task involving a series of images to characterize visual responses. In the natural session (Figure 1B), a human experimenter entered the room containing the arena, offered food to the monkey and left the room. Both sessions were conducted in a large naturalistic arena (Figure 2A) connected through a series of movable partitions to a room containing a touchscreen workstation with gaze tracking capabilities (Figure 2B), as described previously (Jacob et al., 2021a). We performed wireless brain recordings from 256 electrodes implanted into the higher order visual and motor regions (IT, PMv, PFC) of two monkeys (Figure 2C; see Methods).

**Figure 1.**
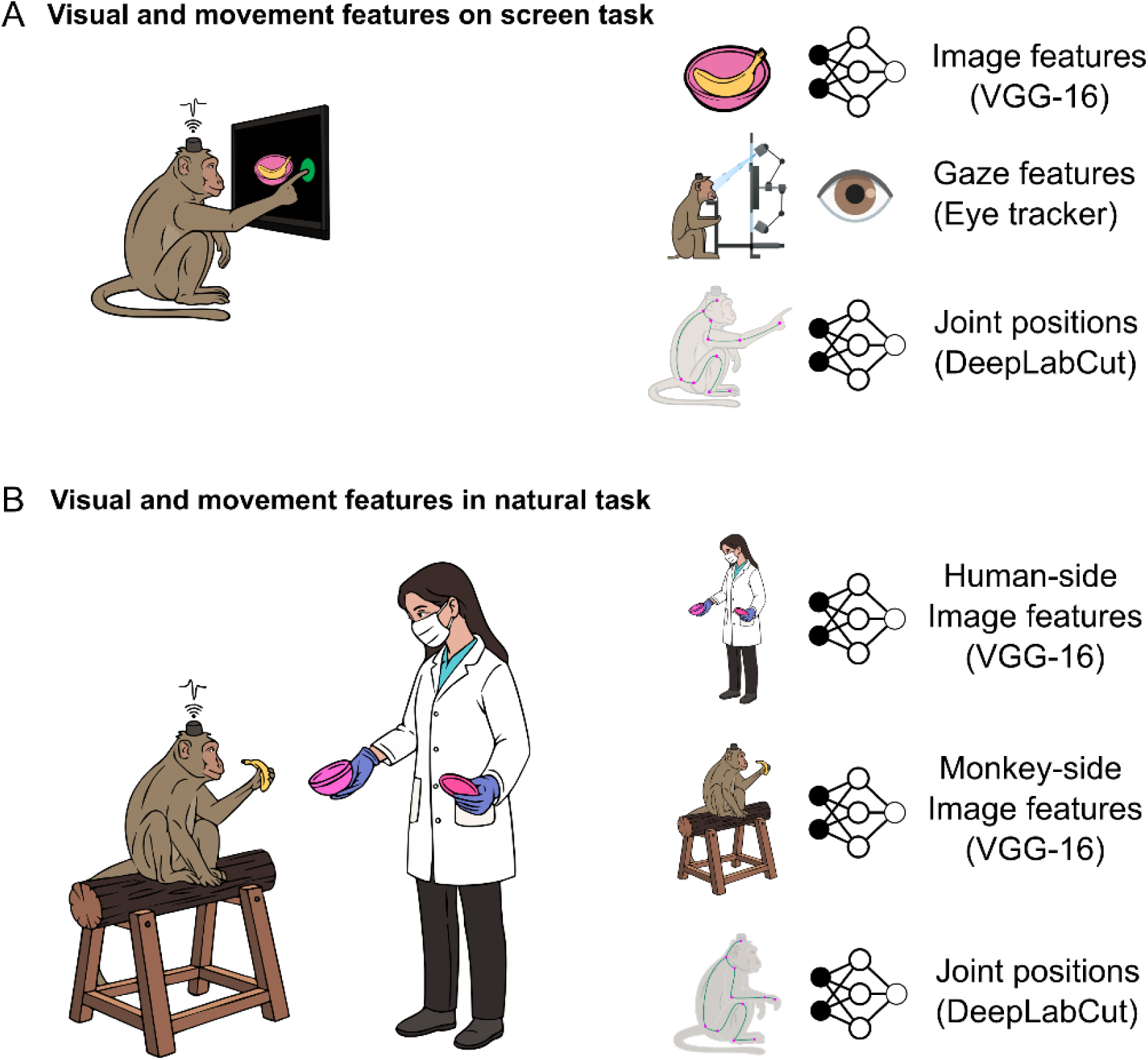
Studying visual and motor information in controlled and natural settings. (A) Schematic of a monkey performing a controlled screen task (*left*), where the monkey had to hold a button on the screen and fixate on a series of images presented at the centre of the screen. During this task, we extracted visual and motor features that might be encoded by neurons in each brain region. These included: image features using a deep network (VGG-16), gaze position (x,y) features from an eye tracker, and joint position (x,y) features using a markerless pose tracking algorithm (DeepLabCut). (B) Schematic of a monkey engaged in a natural task (*left*), in which the monkey was offered various food items by human experimenters. During this task, we extracted visual and motor features that might be encoded by neurons in each brain region. These included human-side image features from a deep network, which we took as the possible visual information seen by the monkey (*right, top*); monkey-side image features from a deep network, which we took as the possible motor features (*right, middle*); and finally, joint positions from an automatic pose tracking algorithm (DeepLabCut; *right, bottom*).

**Figure 2.**
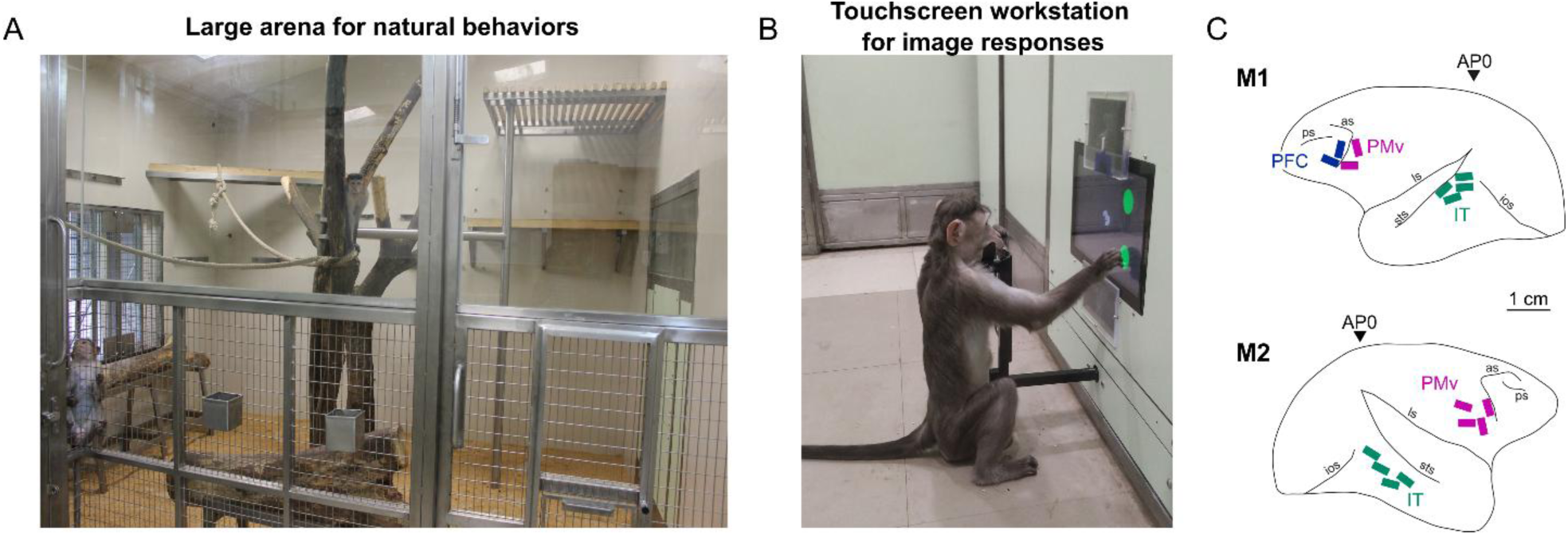
Wireless brain recordings in natural and controlled settings. (A) Brain activity during natural behavior was recorded wirelessly in a large natural arena (3.3 x 4.13 m) enriched with perches, tree trunks, and ropes in which monkeys were group-housed throughout the study. For the natural task, each monkey was separated from its partners using a series of partition doors, one of which is partially visible at the extreme left. (B) Brain activity in controlled settings was obtained using a touchscreen workstation with gaze tracking capability, where each monkey performed a fixation task. The image shows a naïve monkey performing a task. (C) Electrode array locations in each monkey, reconstructed by aligning sulcal landmarks in MRI scans with photographs of the surgically implanted arrays. Monkey M1 had 128 electrodes in IT, 64 in PMv and 64 in PFC. Monkey M2 had 128 electrodes in IT & 128 in PMv. The inverted triangles indicate the stereotaxic zero (AP0). *(This figure is identical to Simon et al, 2026, a previous study using this setup)*

### Dissociation of visual and motor regions in the screen task

Brain activity is often recorded while animals are restrained from making movements. Since movements are often minimized, this approach blinds us from understanding whether movement information reaches or modulates visual activity. However, in our unique touchscreen setup, monkeys were unconstrained and free to make body movements and only had to voluntarily place their head into a chin rest to enable eye tracking and receive juice rewards. Since we knew what they saw on the screen, where they were looking and had simultaneous whole-body video of their movements, we were able to analyze the contribution of joint movements as well as visual information to neural activity in all three brain regions.

We naturally expected that IT neurons would be driven strongly by visual information and that PMv neurons would be driven strongly by motor information – but since these regions are rarely ever recorded from simultaneously, a clear double dissociation has not been observed to our knowledge. We were also additionally interested in characterizing the extent to which motor information predicted IT activity and visual information predicted PMv activity. Ultimately of course, our goal was to investigate the extent to which these findings would hold up in the natural task.

We analyzed neural responses during two fixation tasks. In the first task, monkeys saw images of food, human experimenters as well as play area images, all scaled to be roughly at the size that they would be encountered in the real-world. To avoid large transient effects due to changes in size, these three groups of images (food, experimenter, play area) were presented in separate blocks. To capture the visual features likely to predict neural activity (Figure 3A), we took the penultimate layer activations to the fixation task images from a deep neural network (VGG-16, trained on image categorization) known to predict IT neural responses to natural images (Simonyan and Zisserman, 2014; Jacob et al., 2021b; Katti and Arun, 2022). To investigate whether neural activity is modulated by gaze, we took the (x, y dva) gaze positions from our eye tracker (Figure 3A). To capture the motor features likely to predict neural activity (Figure 3A), we extracted joint positions of the monkey using the automated pose tracking algorithm DeepLabCut (Mathis et al., 2018).

**Figure 3.**
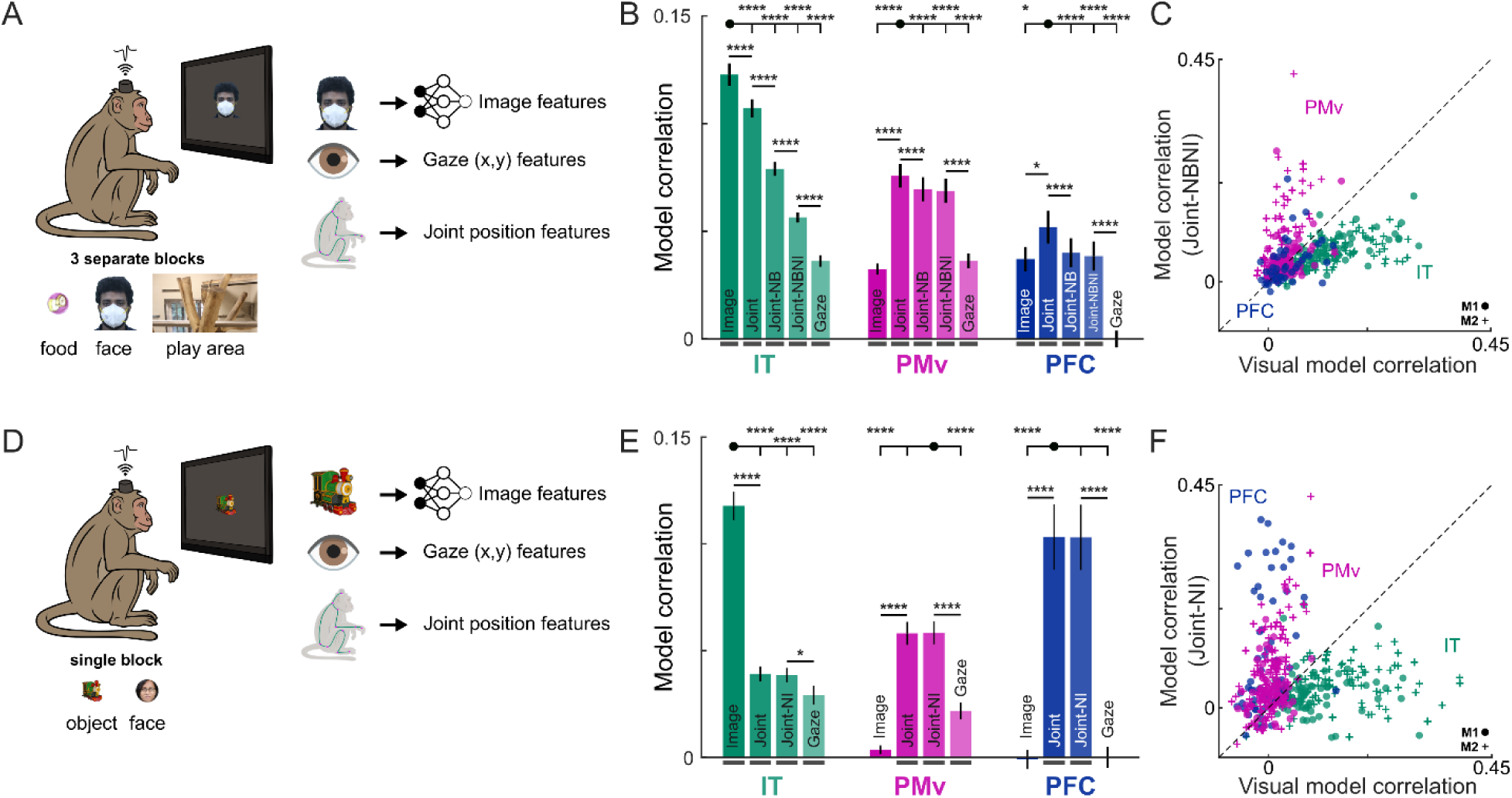
Dissociation of visual and motor regions in two screen tasks. (A) *Left:* Schematic of the fixation task, in which the monkey saw food, face and play area images in separate blocks (depicted to scale below). *[The face image shown is that of one of the authors: D.B.]* *Right:* Schematic of features extracted to predict neural activity: visual features were obtained by passing images into a deep network trained for object categorization (VGG-16); gaze features (x & y eye positions) were obtained from eye tracking; motor features were obtained by extracting the x & y joint positions from the video camera feed of the animals performing the task. (B) Cross-validated model correlation (mean ± s.e.m across neurons) for IT (*green*), PMv (*purple*) and PFC (*blue*). Models whose correlation is significantly different from zero for the neural population are indicated with a *thick horizontal bar*. Asterisks spanning bars represent statistical significance of the corresponding bars, shown for clarity only for consecutive bars and between the highest bar (*black circle*) to all other bars (unmarked pairs are p > 0.05, * is p < 0.05, ** is p < 0.005 etc on a sign-rank test across neurons). *Model abbreviations:* Image: visual features model; Joint: joint position model; Joint-NB: joint positions with block information regressed out; Joint-NBNI: joint positions with block and image onset regressed out; Gaze: gaze features model. (C) Cross-validated correlation of the Joint-NBNI model for each neuron plotted against that of the cross-validated correlation of the visual model, for IT, PMv & PFC neurons (*dots*: monkey M1, *crosses*: monkey M2) (D) Same as (A) but for another fixation task in which object and face images of equal size were presented randomly interleaved within a single block. *[The face image shown is that of one of the authors: S.M.]* (E) Same as (B) but for this fixation task. (F) Same as (C) but for this fixation task, showing a stronger double dissociation.

To predict neural activity, we took the firing rate of each neuron in time bins synchronized to the video feed after the animal began fixating so that visual, motor and gaze features were all available, and built a linear model to predict the firing rate as a weighted sum of either the visual features, gaze positions or the joint positions (see Methods). To evaluate model fits, we performed a 5-fold cross-validation and calculated the correlation between the observed firing rate and the model-predicted firing rate on unseen time bins.

The resulting model correlations are summarized in Figure 3B. As expected, neural responses in IT were predicted best by visual features, compared to joint positions and gaze (Figure 3B). Likewise, neural responses in PMv were best explained by joint positions, and to a smaller degree by gaze and image features (Figure 3B). Finally, neural responses in PFC were best explained by joint features, and next by image features (Figure 3B). All these results are as expected given the known properties of these three brain regions. Note that the model correlations are relatively small because the models are predicting firing rates on individual trials, rather than trial-averaged firing rates as is frequently done for predicting neural responses to images. We confirmed this by comparing model performance trained on trial-averaged firing rates evoked by images with model performance trained on trial-wise firing rates (data not shown).

### Understanding motor contributions to IT neurons

The above results are puzzling, because IT neural responses were explained unexpectedly well by joint positions (Figure 3B). We first wondered whether this could be because of a spurious correlation between the image properties being drastically different across blocks, and the animal shifting his posture systematically across blocks. To resolve this issue, we created a block regressor containing 0/1 values indicating the ongoing block number, regressed out this quantity from each of the joint positions, and used the resulting joint positions to predict neural activity as before (see Methods). The resulting model fits, denoted as “Joint-NB”, showed a larger decrease in model fits for IT neurons compared to PMv neurons (Figure 3B).

Encouraged by this result, we wondered whether joint positions were explaining IT neural responses due to subtle changes in the monkeys’ posture that is somehow time-locked to the onset of images on the screen, in addition to block-related postural changes. To resolve this issue, we created an image regressor containing 0/1 values indicating the presence or absence of any image on the screen, regressed out both the image and block from each of the joint positions, and used the resulting joint positions to predict neural activity as before (see Methods). The resulting model, denoted as “Joint-NBNI”, showed a drastic drop in correlation for IT neurons but only a subtle drop in model fits for PMv neurons (Figure 3B). Thus, IT neural responses are only weakly predicted by joint positions once spurious correlations with image onset and blocks are removed.

To visualize the dissociation between visual and motor features driving neural activity in the three brain regions, we plotted the joint model correlation (Joint-NBNI, after regressing out image & block) against the visual model correlation (Figure 3C). This revealed a striking double dissociation, whereby most IT neurons are predicted better by visual features but not motor features, most PMv neurons were predicted better by joint positions compared to visual features, and PFC neurons were predicted equally well by visual and motor features.

### Markerless pose tracking is biased by nearby image features

The above findings raise the intriguing possibility that joint positions change subtly across blocks and with image onset. This could be due to monkeys actually shifting their posture, but a simpler and more worrying possibility is that automatic pose tracking is biased by the presence of nearby image onset such as the images appearing on the screen. To investigate these possibilities, we first confirmed that block and image onset can indeed be decoded from the monkeys’ joints (Section S1). We then noted that joints closer to the screen tended to carry more information, suggesting that automatic pose tracking is indeed affected by image onset and by image size. To comprehensively confirm this possibility, we created dummy videos containing the monkey in exactly the same pose, but with or without a white rectangle overlaid as an image on the touchscreen. This revealed a systematic change in estimated wrist and palm position, confirming that markerless pose tracking is indeed modulated by image onset (Section S1). This in turn confirms that the joint position model was indeed explaining visual responses due to this artefact of the pose tracking algorithm. Thus, joint positions estimated using markerless pose tracking can be systematically biased by nearby image features and must therefore be used with caution, particularly in conjunction with visual feature analyses.

### Dissociation between visual and motor regions in another screen task

The above results show that, in a fixation task where images of different sizes are presented in blocks, a clear double dissociation between visual and motor features can be obtained between IT & PMv, but only after accounting for variations in joint positions due to image onset or block variations. We found that the correlation between IT neural activity and joint positions are most likely spurious: they could arise from real postural changes made by the monkey or for that matter, arise from joint position information reaching IT neurons, and we were able to remove these correlations by factoring out image onset information from the joint positions. To investigate this issue further, we repeated these analyses on another fixation task in which monkeys saw images equated in size (8 objects and 8 faces equated in size; see Methods) and presented randomly interleaved within a single block (see Methods; Figure 3D).

The resulting model fits revealed an even more striking double dissociation: IT neurons were predicted much better by visual features compared to motor features; PMv neurons were predicted much better by motor compared to visual features and PFC neurons were predicted largely by motor compared to visual features (Figure 3E). In this experiment, removing image onset information from the joint positions did not change model predictions (Joint-NI vs Joint model; Figure 3E). To visualize the double dissociation between visual and motor information, we plotted the cross-validated correlation of the Joint-NI model against that of the visual model (Figure 3F). This revealed a striking double dissociation at the single-neuron level, whereby most IT neurons were predicted better by visual features, most PMv neurons were predicted better by joint positions, and PFC neurons were predicted better by joint positions (Figure 6F).

Taken together the above results show that high-level visual and motor regions are doubly dissociated in well-controlled screen tasks.

### Dissociation of visual and motor regions in the natural task

The above sections show that visual and motor regions show a clear double dissociation in well-controlled screen tasks. We therefore wondered whether this dissociation would be present in the natural task. To investigate these possibilities, we took the firing rate of each neuron during the natural task in time bins synchronized to the video feed. Since our natural arena contained a grill partition separating humans and monkeys, we realized that the video recordings from each side of the partition could be used to extract visual and motor features (Figure 4A).

**Figure 4.**
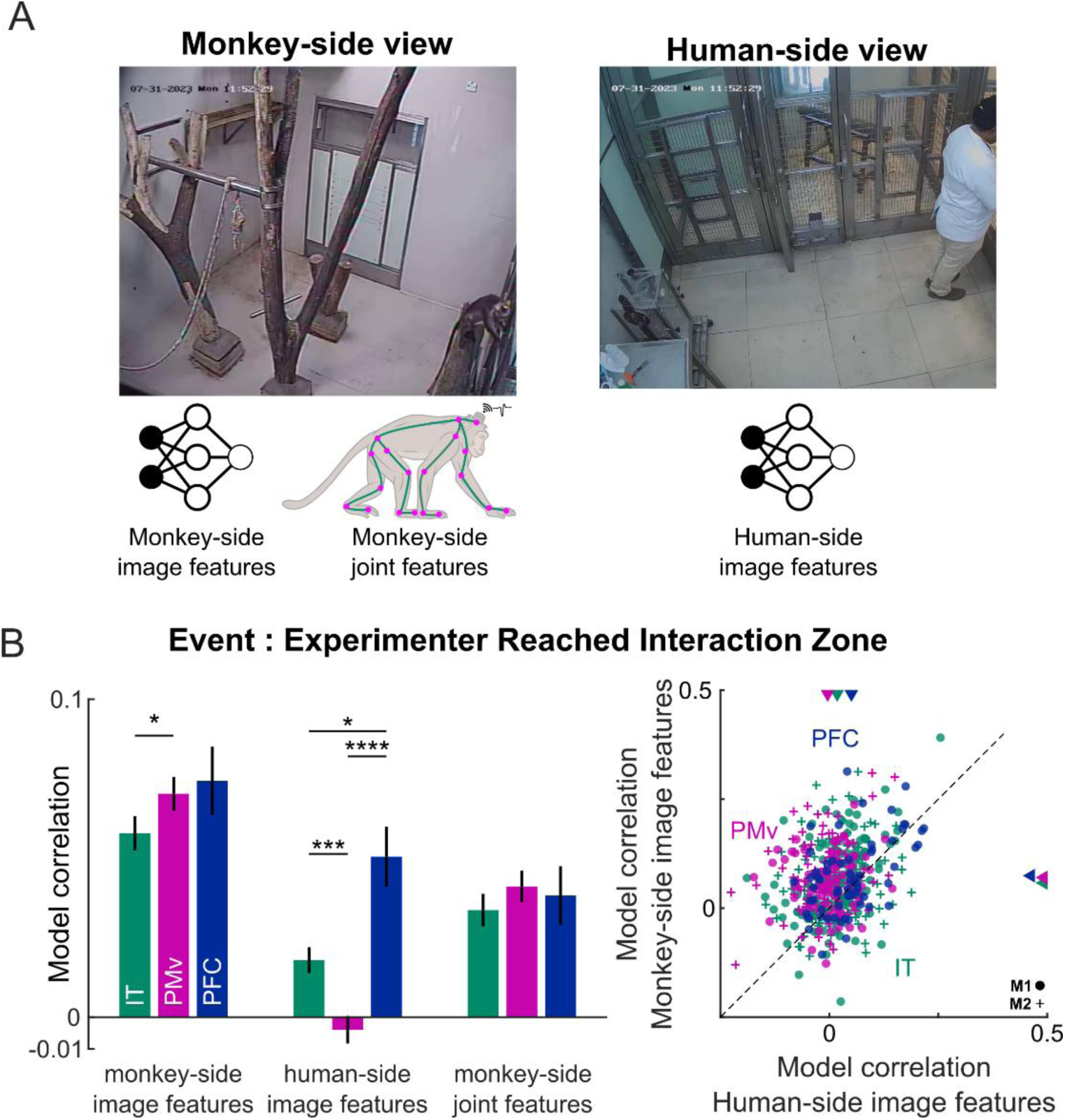
Dissociation of visual and motor regions in the natural task. (A) Example frames showing the monkey-side and human-side views during the natural task. The human in the frame is one of the authors (D.B.). For the monkey-side view, we extracted visual features by passing frames through a deep network and extracted joint positions using automated pose tracking. For the human-side view, we extracted visual features by passing the frames through a deep network. (B) *Left:* Cross-validated model correlation (mean ± s.e.m across neurons) for IT (*green*), PMv (purple) and PFC (blue) for monkey-side image features, human-side image features and monkey-side joint features. Model correlations were evaluated in a 1 s window around the event when experimenter reached the interaction zone. Asterisks represent statistical significance evaluated using a sign-rank test comparing the model correlations across neurons (* is p < 0.05, ** is p < 0.005 etc). Note that in this plot, model correlations are grouped by models and compared between brain regions (instead of between models in each brain region, as in figure 3B, E), because we wanted to compare amount of information encoded by visual vs motor brain regions during unrestrained natural behaviour. *Right.* Cross-validated model correlation for monkey-side image features plotted against the cross-validated model correlation for human-side image features. IT and PMv show a weak dissociation during this event.

We therefore extracted three possible predictors of neural activity: visual features from the monkey-side view obtained by passing these images through a deep network (VGG-16, trained on ImageNet) as before, joint positions extracted by automated pose tracking from the monkey side view, and visual features extracted from the human side. While the human-side image features contain purely visual information, the caveat with the monkey-side image features is that the image contains both visual features of objects seen by the monkey as well as motor features when the monkey is present in the frame. For each neuron, we predicted the firing rate across the entire experiment as a weighted sum of the monkey-side image features, human-side image features and monkey-side joint features. For each model, we calculated the model correlation by using a leave-one-condition-out cross-validation, where a condition is an experimenter-food pair in the natural interaction task. Since the natural task contained several distinct events, we evaluated this model correlation in a window-0.5 to 0.5 s of each event.

The resulting model correlations are shown in Figure 4B, for the event when the human experimenter reached the interaction zone to offer food to the monkey. We found that monkey-side image features explain PMv responses better than IT, whereas human-side image features explain IT responses better than PMv (Figure 4B). However this dissociation (Figure 4B, right) was nowhere as clear as observed in the screen tasks (Figure 3C, 3F), presumably because both monkey-side and human-side events are correlated: for instance, as the experimenter moved in the human-side view towards the monkey, the monkey also moved in matched fashion to receive the food, introducing a natural correlation between the visual features on both sides. Indeed, repeating this analysis for other events in the natural task revealed a rich, dynamic picture in which some events show a clear dissociation whereas others did not (Section S2). Importantly, visual and motor features were least correlated for events that showed a clear dissociation (Section S2).

We conclude that visual and motor features are dissociated in the natural task only at specific events where presumably these features are themselves relatively uncorrelated.

## DISCUSSION

Here, we investigated the degree to which visual and motor information is encoded in high-level visual and motor regions during controlled tasks and during natural behaviors. To address these issues, we performed wireless recordings in high-level sensory and motor regions (IT, PMv, PFC) of freely moving monkeys performing a natural task in a large natural arena and also performing controlled tasks on a touchscreen workstation. Our main finding is that high-level visual and motor regions are doubly dissociated in screen tasks but only during specific events in the natural task. Below we discuss this finding in the context of the existing literature.

### Dissociation of visual and motor regions during natural behavior

We have found a striking double dissociation between IT & PMv in well-controlled screen tasks: neural responses in IT were predicted better using visual but not motor features, whereas PMv responses were explained better using motor but not visual features. While this finding as such is not unexpected since IT is a visual region and PMv is a motor region, this double dissociation has not been directly demonstrated previously under matched conditions. This is important because IT neurons are usually tested with a large diversity of stimuli albeit while restricting monkeys’ movements, and PMv neurons are usually tested using a large diversity of movements while presenting highly impoverished stimuli, leaving open the possibility that these two regions might behave differently when tested with diverse visual stimuli and motor movements. Reassuringly, this was not the case. Likewise, there has been some recent debate regarding whether movements drive neural responses in the visual cortex (Kang et al., 2025). Our results are concordant with the recent finding that spontaneous movements have little impact on primate visual cortex (Talluri et al., 2023). However, we demonstrate this result in a more general and natural setting since our monkeys were unrestrained and had many more degrees of freedom of movement. Yet we found very little influence of movement, that too in IT cortex which is a higher-level visual region that is anatomically connected to prefrontal and motor regions (Conway, 2018; Ayzenberg and Behrmann, 2022). Our results also extend these ideas to the premotor cortex by showing that PMv neurons are strongly driven by joint movements as expected, but importantly, are minimally driven by visual features.

### Limitations of this study

We have found that the strong dissociation between IT & PMv in well-controlled screen tasks is far weaker in natural tasks, and can be seen only at specific events. However, this weaker dissociation cannot be interpreted as proof of greater mixing of visual and motor information during natural behaviors, because many simpler explanations will need to be tested and rejected.

First, visual features used for predicting neural activity in our analysis (monkey-side and human-side image features) may not have captured the exact visual information seen by our monkeys. Indeed, this is a major limitation of our study, since it was technically challenging to record the monkeys’ gaze and their visual inputs. Fortunately there are exciting attempts in that direction, with either cameras directly implanted on the head (Singh et al., 2025), or having arrays of cameras that can be used to reconstruct any viewpoint (Voloh et al., 2023). Second, even the joint movements too were only two-dimensional joint positions which might have sufficed for the screen tasks since movements were largely planar but might not have captured the 3-dimensional trajectories of joints during natural behavior. Finally, visual and motor features in the natural task might have been correlated – for instance, the monkey moved to receive the food as the experimenter approached, introducing a natural correlation between its visual input and motor movements. Such correlated behavior can result in neural activity in visual cortex becoming predictable by joint movements. We have also found that automated pose tracking algorithms can introduce spurious correlations because their estimates of joint positions near the screen can be biased by images on the screen (Section S2). These findings taken together underscore the need for caution in interpreting any apparent mixing of information observed during natural behaviors.

### Conclusions

Taken together, our results reconfirm the traditional view that sensory and motor regions indeed encode distinct types of information, and that these findings hold even when these regions are tested in unrestrained monkeys under natural settings with a far larger range of movements.

## METHODS

All animal experiments were performed according to a protocol approved by the Institutional Animal Ethics Committee of the Indian Institute of Science, Bangalore (CAF/Ethics/750/2020) and by the Committee for the Purpose of Control and Supervision of Experiments on Animals, Government of India (V-11011(3)/15/2020-CPCSEA-DADF). Many experimental procedures are common to previously reported studies from our laboratory (Ratan Murty and Arun, 2015; Jacob et al., 2021a), and summarized briefly below.

### Neurophysiology & wireless recordings

We implanted two adult bonnet macaque monkeys (Ma*caca radiata,* M1: *Co* & M2: *Di*, both aged 9.5 years) with Floating Microelectrode Arrays (FMA) from Microprobes for Life Science using standard neurophysiological procedures reported previously (Ratan Murty and Arun, 2015) adapted to procedures required for chronic wireless recordings as described recently (Simon et al., 2026)

### Touchscreen workstation

Monkeys performed the screen-based tasks on a touchscreen workstation with accurate eye-tracking capabilities as reported previously (Jacob et al., 2021a; Simon et al., 2026).

### Alignment of video recordings to neural data

Although video cameras at the touchscreen workstation were fully synchronized with the neural data acquisition system, the video cameras in the natural arena were not synchronized in this manner. To achieve alignment, we created a synchronization task as described previously (Simon et al., 2026). Briefly, full screen images were presented at precise times on the touchscreen, which were captured by the synchronized and unsynchronized cameras and subsequently used for synchronization.

### Natural task

To study natural vision, we devised a simple behavioral task. On each trial, a human experimenter entered the natural arena with a covered box of food, approached the interaction zone and opened the lid of the box to reveal the food, upon which the monkey took the food from the box. In all, one of three types of food (pieces of banana, carrot or raisin) was offered by one of two experimenters (the authors D.B & S.M). The food box was offered so that it was close to the monkeys’ right (dominant) hand in the first 24 trials and to the left hand in the next 12 trials. The sequence of trials was randomly chosen using a computer with the constraint that consecutive trials would not have the same food or experimenter. There were 37 trials for monkey M1 and 36 trials for monkey M2.

We manually annotated the videos which contained the clearest view of the monkey-human interaction for the times of key events during each trial. The key events were chosen after carefully reviewing the entire session video to identify distinct, repeating and well-defined events that occurred across trials. The annotated events were: Experimenter Entered Room (EER), Experimenter Left Room (ELR), Experimenter Show Box (ESB), Experimenter Hide Box (EHB), Experimenter Reached Interaction Zone (ERIZ), Experimenter Left Interaction Zone (ELIZ), Experimenter Offered Food (EOF), Experimenter Offered Hand (EOH), Experimenter Withdrew Hand (EWH), Monkey Start Reach (MSR), Monkey End Reach (MER), Monkey Start Withdrew Hand (MSWH), Monkey Stop Withdrew Hand (MEWH), Monkey Held Food (MHF), Monkey Held Hand (MHH), Monkey Held Grill (MHG), Monkey Released Grill (MRG), Monkey Face Experimenter Entered Room (MFEER), Monkey Look At Food (MLF), Monkey Look At Empty Box (MLEB), Monkey Look At Experimenter (MLE), Monkey Look At Experimenter Hand (MLEH), Monkey Start Climb Log (MSCL), Monkey End Climb Log (MECL), Monkey Reached Interaction Zone (MRIZ), Monkey Left Interaction Zone (MLIZ), Monkey Startled (MS).

### Touchscreen tasks

Before the natural task recordings, both monkeys performed two touchscreen-based tasks: a visual task and a motor task.

*Eye calibration block.* Each task started with a block for calibrating eye position, as detailed previously (Simon et al., 2026). Briefly, the monkey had to complete two correct calibration trials by fixating successively on four gray circles (radius 0.5°) that appeared consecutively and randomly at four corners of a square of side 16°. Trial was considered correct if monkey maintained fixation in each of the circles within a certain window for 0.3s. The eye data from these trials was then used to learn a linear transformation between the raw eye signal voltage and the visual angles of each dot on the screen. This transformation was later used for all trials to calibrate the eye signal of monkeys post-hoc.

*Visual task.* Each monkey performed a passive fixation task as described previously (Simon et al., 2026). Each trial began with a blue circle serving as a hold button (4° radius), positioned 28° to the right from the screen centre. The monkey had 10 s to touch this button. To accommodate differences in touching behaviour, touches were accepted within 8° of the button for M1 and within 2° for M2. Touching the button brought up a yellow fixation dot (0.1° radius) at the screen centre. The monkey then had 0.5 s to bring its gaze within a 10° radius of this dot. Once fixation had been maintained for 0.3 s, a sequence of eight images were shown in succession, each lasting 0.2 s and separated by a 0.2 s blank interval. Throughout this sequence, the monkey was required to maintain fixation within the same window and received a juice reward upon successful maintenance of gaze. Releasing the hold button or breaking fixation terminated the trial and this trial was repeated after a random number of intervening trials. We initially chose a relatively broad fixation window because fixation accuracy was uncertain. Subsequent analysis, however, showed that both monkeys held their gaze accurately throughout the trials. The within-trial standard deviations of horizontal and vertical eye position, averaged across trials, were 1.05° and 0.99° for M1, and 0.71° and 0.91° for M2, respectively. Mean eye-position jitter across trials was 0.37° horizontally and 1.3° vertically for M1, and 0.58° horizontally and 0.80° vertically for M2.

The main fixation task (Simon et al., 2026) comprised three image sets, presented in separate blocks. The first contained food-related images chosen to approximate themonkeys’ views during the natural task. These included a food box containing banana, carrot or raisin pieces, photographed from five viewpoints for each food type (3 × 5 = 15 images), along with the empty box photographed from the same five viewpoints (5 images). A further 9 images showed the three food types on the floor from three viewpoints each (3 × 3 = 9), alongside three images of the empty floor, giving 32 images in this set. The second set depicted three human experimenters: E1 and E2, who would later take part in the natural task, and E3, who served as a control. Each experimenter was photographed from five viewpoints, both with and without a face mask, yielding 30 images (3 experimenters × 2 mask conditions × 5 views). The third set comprised 18 images sampled from the natural arena. Image sizes were selected to approximate their likely apparent sizes during the natural task: food images measured 7.8° along their longest dimension, face images subtended 22.2°, and arena images measured 40.8° along their longest dimension. Across the experiment, each of the 80 unique images appeared 16 times over 160 correct trials. Images were presented in random order within each trial, subject to restrictions on consecutive presentations. In the food and face blocks, successive images could not repeat the same viewpoint, food type or face. Arena images were randomly interleaved across categories, with no two consecutive images belonging to the same category, such as perches or logs.

### Extraction of joint position model features using markerless pose tracking

We used a markerless automatic pose tracking software, DeepLabCut v2.3.8 (Mathis et al., 2018) to estimate the joint positions to be used for predicting neural responses. We identified 19 canonical joints in each monkey: head, neck, hip and both left & right shoulders, elbows, wrists, palms, side-hips, knees, ankles and feet. In addition, we also annotated the wireless logger as an additional feature since it was a salient and easily tracked feature for both monkeys. Since absolute (x, y) positions of each joint in the video image frame are entirely arbitrary, we calculated the (x, y) positions of each joint relative to the wireless logger to obtain the position of the joint relative to the body of the monkey. In practice this does not change our results since it only amounts to changing a constant term in our neural prediction models.

To train the DeepLabCut algorithm on each camera output, we manually annotated each joint position in 99 randomly chosen frames (random 27 frames from M1 and random 72 frames from M2 session) across both monkeys from videos of them performing the right-hand motor task as captured from our sideview camera. Based on DeepLabCut documentation and our experience, this much training data was sufficient to get ceiling performance for unsupervised training within the DeepLabCut environment. We then trained the DeepLabCut software using its prepackaged resnet152 network for training and evaluation purposes using these 99 frames as training data. We stopped model training after 200,000 iterations, at which point performance had saturated to a training error of 2.36 pixels, and a test error of 5.22 pixels with a likelihood cutoff for each joint set at 0.6 during network evaluation.

We then used this model to estimate joint positions of each monkey’s videos while it performed the visual task. In the model output, we retained only the estimated joint positions with average likelihood > 0.5 and set the remaining values to NaN (not a number). This likelihood threshold was chosen by inspecting performance of tracking of each of the joint positions on a separate held-out manually-annotated dataset for each camera view (thus chosen joints for the side vs top view cameras are different; data not shown). This step ensured that only high-quality joint positions were used for predicting neural responses. For the joint position model, we retained the (x, y) joint positions of all valid joints from the time of the first image onset in the trial to the end of the last stimulus off time, which was the period during which we had valid gaze data. These joint positions were concatenated across all frames and all trials for predicting the neural response (as detailed below).

### Extraction of visual model features using pretrained deep networks (VGG-16)

We used a pretrained deep convolutional neural network (VGG-16) trained on object categorization to extract visual features to be used for predicting neural responses. In the visual task, we generated 60 fps video of what the monkeys saw during the fixation task using the same time window of the trial as before (using the *mlplayer* function, MonkeyLogic). We then downsampled these videos to 30 fps to align them to the frame rate of the videos used for joint position estimation. We then cropped a rectangle (40.8° x 22.2°, which is the max across all images in the entire experiment along their longer dimensions) to capture the region of the screen visible to the monkey during passive fixation task. We then passed each of these cropped frames through the pretrained VGG-16 network pre-trained on categorization on the ImageNet database (*vgg16* function, Deep Learning Toolbox, MATLAB 2025) and collected its activation to each of the frames from the ‘fc8’ layer (last fully connected layer), which yielded a 1000 x 1 vector for each frame (other layers such as ‘fc6’ or ‘conv5’ gave us qualitatively similar results). We concatenated these features across all frames and all trials for predicting the neural response (as detailed below).

### Extraction of features for gaze model

We extracted the x & y positions from the calibrated eye position data during the same part of the trial as before. We down-sampled and aligned the eye data to the framerate of the joint movements (30 fps), which gave us two values per frame. These 2 gaze features were concatenated across all frames and all trials for predicting the neural response (as detailed below).

### Linear regression for joint, visual and gaze models

To predict the neural response, we calculated the firing rate in time bins corresponding to the joint position video frame rate (30 fps), and concatenated these firing rates across time bins and trials to obtain a single vector. To reduce redundancy, we performed a dimensionality reduction for each feature bank (visual or motor) using principal component analysis and retained the components that explained 90% of the variance. For each model, we solved the linear regression ***y*** = ***Xb*** where ***y*** is a nframes x 1 vector of firing rates (nframes being the number of frames), ***X*** is a feature matrix of dimension nframes x (nfeatures + 1), containing the features of each frame along with a constant term and ***b*** is a (nfeatures + 1) x 1 vector containing the unknown weights of summation required to predict the firing rate. The final size of the regression matrices for each model were: 15520 x 17 for joint positions, 15220 x 12 for visual features and 15220 x 3 for gaze features (same for M1 & M2 since both had equal number of correct trials, and thus equal number of frames from first image on to last image off per trial).

To evaluate each model, we performed a 5-fold cross-validation as before: we separated all the frames into 5 non-overlapping groups, and trained the model on 4 groups at a time and calculated model predictions for the held-out group. We concatenated the model output across each held-out group to obtain model predictions for all frames together, and calculated the correlation with the observed firing rates. Varying these choices yielded qualitatively similar results.

### Joint-NB & Joint-NBNI models

To remove possible covariations of joint positions with block and image onset, we modified the joint position features as follows. To create the regression matrix for the Joint-NB model, we created a block ID vector containing two values [isblock1 isblock2] for each frame, where isblock1 is 1 for block 1 frames and 0 for others, and isblock2 is 1 for block 2 frames and 0 otherwise. Block 3 frames are then indicated by both values being zero. For each joint position in the original joint position matrix, we solved a linear regression of the form ***j*** = ***Bw*** where ***j*** is a nframes x 1 vector of estimated joint positions and ***B*** is a nframes x 2 vector containing the block ID vectors and ***w*** is a vector of unknown weights that we solved using linear regression. We then created a new joint position vector by taking ***j_new_*** = ***j*** − ***Bw*** which now has reduced or no dependence on block information.

For the Joint-NBNI model, we included an additional column into the block ID matrix which indicated whether an image was present (1) or absent (0) in that frame, and repeated the above procedure.

### Visual and motor features in the natural task (Figure 4)

In the natural task, we had video recordings in which the human experimenter was always on one side of the grill and the monkey was on the other side. We therefore created three models. In the human-side visual features model, we passed the human side of the video frame where the monkey was not visible, into the same pretrained deep network as above (VGG-16), and extracted the activations from the fc8 fully connected layer. In the monkey-side visual features model, we did likewise for the monkey-side of the video frame which contained the monkey and the natural arena features but not the human. In the monkey-side joint position model, we estimated joint positions of the monkey using DeepLabCut (by annotating total 148 frames for monkey-side camera and training the algorithm in a similar fashion as before). For each feature bank, we reduced dimensionality using PCA to retain 99% of the variance.

For each event, we considered a-0.5 to 0.5 s window relative to the event time, which yielded 312 frames for regression, and we calculated the response of each neuron during these same frames, and performed a linear regression as before. To evaluate these models, we performed a leave-one-condition-out cross-validation: we removed all frames of a given condition (experimenter-food type combination) and trained each model and evaluated its predictions by concatenating model predictions as before. These fits are reported in Figure 4 & Figure S3/ Section S2.

## ACKNOWLEDGEMENTS

We thank Dr. Ashesh Dhawale for valuable discussions, Dr. Tim Blanche of White-Matter LLC for technical support with wireless recordings, Dr. Sebastian Chandu for assistance with implant surgeries, Dr. Sumedh Shastry for help with physiology during surgery and post-surgical care, Dr. Sumedh Shastry & Mr. V. Ramesh for primate facility management and animal care and Mr. Ravi & Mr. Ashok for animal care.

## AUTHOR CONTRIBUTIONS

Shubhankar Saha: Recording setup, Experiment Design, Data Analysis, Manuscript writing and editing. Specific contributions: Developed hardware & software for wireless brain recordings, trained monkeys, planned and conducted implant surgeries, performed automated pose tracking, performed data analysis, wrote manuscript, provided feedback on the manuscript drafts.

Sini Simon: Recording setup, Manuscript editing. Specific contributions: Developed hardware & software for wireless brain recordings, tested electrodes, trained monkeys, planned and conducted implant surgeries, participated in troubleshooting recordings, provided feedback on manuscript drafts, incorporated feedback from all authors on earlier manuscript versions.

Surbhi Munda: Recording setup, Experiment Design, Data Collection & Curation. Specific contributions: Developed hardware & software for wireless brain recordings, trained monkeys, planned and conducted implant surgeries, participated in troubleshooting recordings, designed experiments, collected data, curated data, helped with data visualization, provided feedback on the manuscript drafts.

Thomas Cherian: Recording setup, Experiment Design. Specific contributions: Developed hardware & template software codes for recording, tested electrodes and implant housing, planned and conducted implant surgeries, participated in troubleshooting recordings, and designed motor tasks.

Georgin Jacob: Recording setup. Specific contributions: Developed hardware & template software codes for recording, developed software for automated spike detection and sorting, tested electrodes and implant housing, planned and conducted implant surgeries, participated in troubleshooting recordings, and provided feedback on manuscript drafts.

Jhilik Das: Recording setup, Manuscript editing. Specific contributions: Developed hardware & software codes for data curation and analysis, tested electrode and implant housing, snout trained monkeys, planned and conducted implant surgeries, participated in troubleshooting recording setup, and provided feedback on manuscript drafts.

Debdyuti Bhadra: Recording setup, Experiment Design, Data collection, annotation, visualization. Specific contributions: Designed experiments, collected data, helped with hardware and software updates for wireless brain recordings, planned and conducted implant surgeries, annotated videos, assisted with automated pose tracking, provided feedback on manuscript drafts.

## DATA AVAILABILITY

All data and code required to reproduce the results are publicly available at https://osf.io/j73qp

## SUPPLEMENTARY MATERIAL

## SECTION S1. ANALYSIS OF MARKERLESS POSE TRACKING

In monkeys performing a fixation task involving images of different size presented in blocks, we observed that neural activity in IT was explained surprisingly well by joint positions. This could be because monkeys were shifting their posture systematically across blocks, and these shifts might have correlated with the changes in IT firing rates due to varying image size. We investigated this issue in the main text by regressing out block and image onset information from the joint positions, which greatly reduced the ability of joint positions to predict IT firing rates.

The above results could arise if there was subtle changes in the monkeys’ posture across blocks, or due to artefacts in automated pose tracking introduced due to images appearing near the monkeys’ hand which was touching the screen. We investigate these issues extensively below.

### Decoding of block information from joint positions

We first started by plotting the average estimated joint position of monkey M1 and M2 in the three blocks (Figure S1A). Although we did not observe any visible difference in the monkeys’ posture, the estimated joint positions were slightly different in the three blocks (Figure S1A). Interestingly, this effect was slightly larger for joints closer to the touchscreen. This could affect joints that are further away also, since the pose tracking algorithm might assume connections between joints.

To confirm whether block information can indeed be decoded from each joint, we trained a linear decoder on the (x,y) positions of each joint to categorize the block identity (3-way, since there were 3 blocks), and evaluated the performance of this decoder using 5-fold cross-validation. This revealed that block information could be reliably decoded by nearly all joints in both cameras and in both monkeys (Figure S1B).

### Decoding of image onset from joint positions

The above findings suggest that block information is present in joint positions, reflecting either a subtle change in posture or biasing of pose tracking due to images on the screen. To test the latter possibility, we performed an additional analysis to decode the presence of an image on the screen. If image on/off information can be decoded reliably from joint position, it would indicate that pose tracking is biased by the image onset since it is highly unlikely that monkeys would be shifting their pose in a manner time-locked to image onset.

We therefore trained a linear decoder on the (x,y) positions of each joint across frames to decode whether an image is present on the screen or not, and evaluated the performance of this decoder using 5-fold cross-validation. This revealed that image onset could be reliably decoded from the head, neck, right shoulder, right wrist and palm in both monkeys (Figure S1C).

These results further confirm the possibility that pose tracking is systematically affected by nearby image features, thereby picking up information about block and image onset.

### Direct proof that pose tracking is biased by nearby images

The above results still do not rule out completely the possibility that monkeys were making subtle adjustments to their joint positions during the experiment. To completely rule out this possibility, we created two dummy video files with a single frame containing a frozen view of one monkey, such that one video contained a blank screen, and the other video contained a white square of 80 x 80 pixels on the touchscreen (Figure S2A). We then passed each of these videos through our pose model (trained on the same camera feed, check Methods on Pose tracking using DeepLabCut) and collected the joint positions.

The resulting joint positions are shown in Figure S2A. When we plotted the total change in joint position (pixels) between both these conditions, we found that the right wrist and the right palm, which are closest to the white square, were deviated in their position (Figure S2B).

These results provide direct proof that markerless pose tracking algorithms are biased by nearby image features. Thus, these algorithms must be used with great caution particularly when analysing them in conjunction with visual features.

**Video S1. Demonstration of how automated pose tracking can be biased by nearby image features (Video S1).**

The video shows joint positions of monkey M2 estimated using DeepLabCut overlaid on its body as it performs a fixation task with small, medium and large-sized images.

---

**Figure S1.**
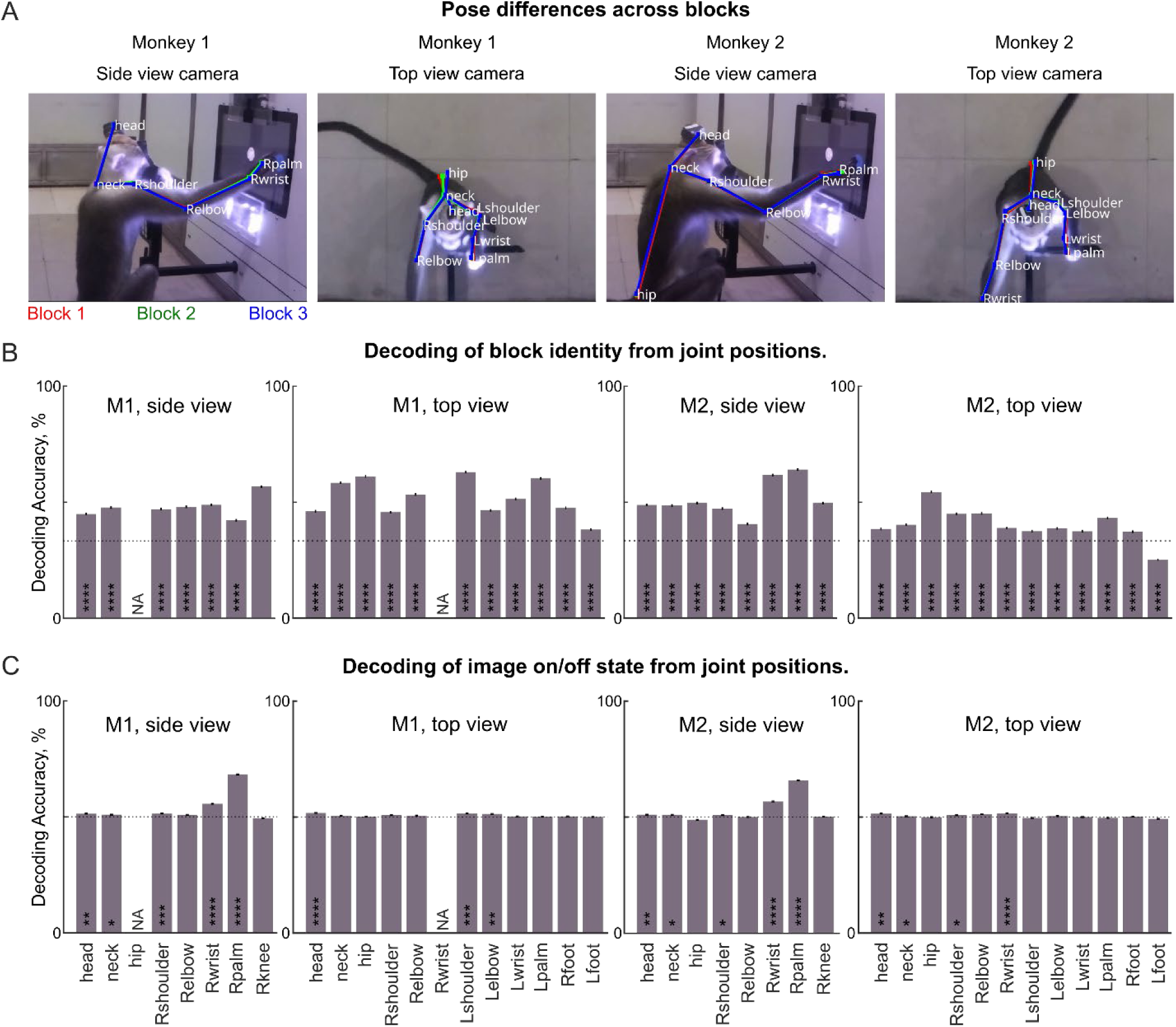
Decoding of block and image onset from estimated joint positions (A) Average joint positions of Monkey M1 & M2 estimated from the side view & top view cameras for Block 1 (red), 2 (green) and 3 (blue) in the fixation task, overlaid on an example video frame. Joint positions are shown for reliably tracked joints in both monkeys and were averaged across all correct trials from the first image onset to the last image offset. Note that there are slight differences in joint positions across blocks. (B) Cross-validated decoder accuracy of a decoder trained on (x,y) pixel position of each joint to decode block identity (1/2/3) for each view and monkey. Asterisks inside each bar indicate statistically significant decoding (**** indicates p < 0.00005, chi-squared test). “NA” indicates joints that are not reliably tracked. (C) Cross-validated decoder accuracy of a decoder trained on (x,y) pixel position of each joint to decode image presence in the frame (0 = image absent, 1 = image present) for each view and monkey.

**Figure S2.**
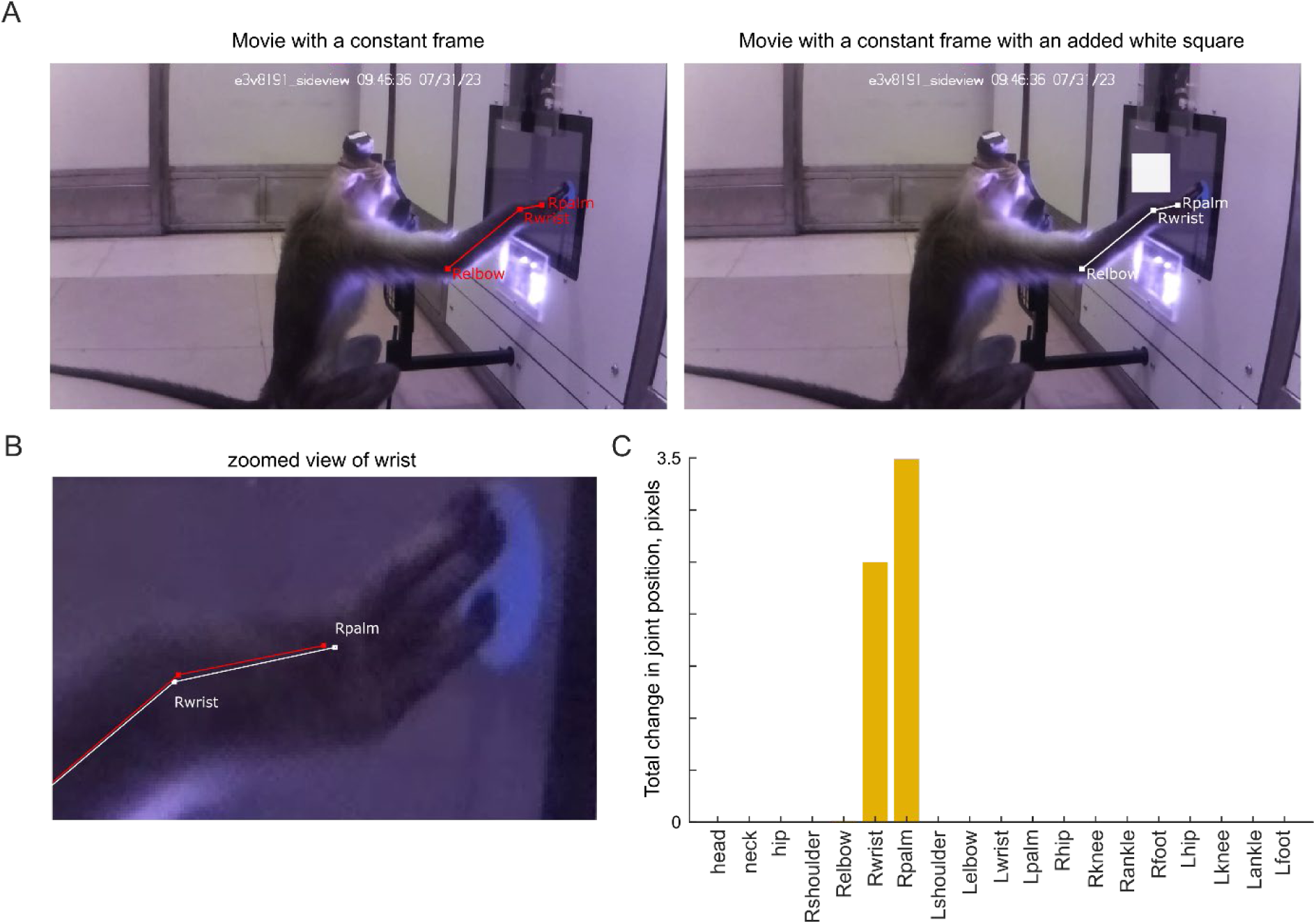
Direct proof that pose tracking is biased by nearby image content (A) *Left:* Example frame from Monkey M2 used to create a dummy video with no monkey movement, containing a blank screen. *Right:* Example frame from a second dummy video created by adding a white square to the dummy video with the blank screen. (B) Estimated joint position from the blank screen video (*white*) and the white square video (*red*), showing a slight change where there should have been none (since the monkeys’ joints have not moved). (C) Total change in estimated joint position (in pixels) for each joint. It can be seen that only the right wrist and palm are affected, which are the two joints closest to the white square. Thus, automated pose tracking is affected by nearby visual features.

**Figure S3.**
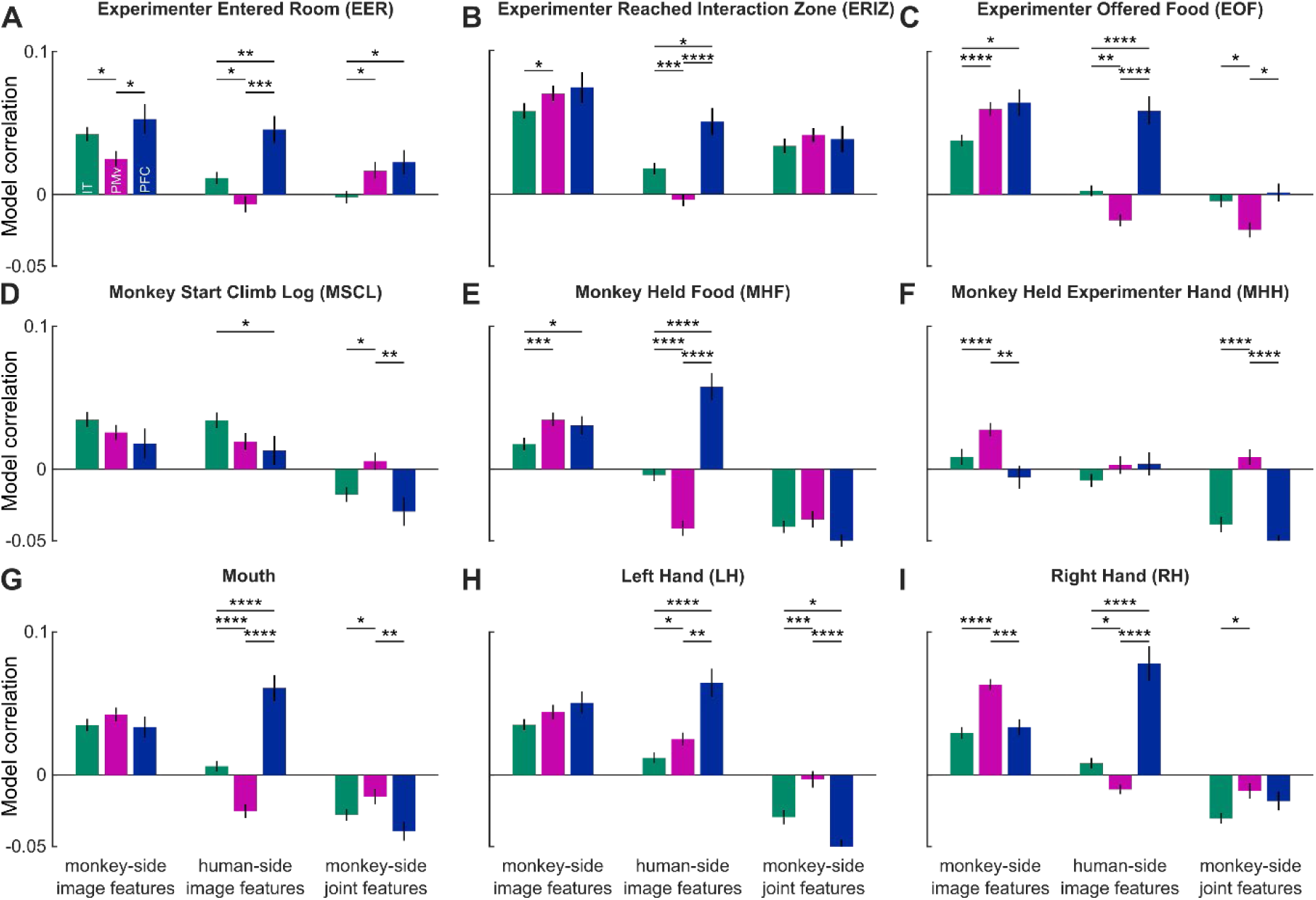
Dissociation of visual and motor regions in the natural task. Each plot shows the cross-validated model correlation (mean ± s.e.m across neurons) between predicted and observed firing rate in a window-0.5 to 0.5 s relative to the event marked in the title, for IT (*green*), PMv (*purple*) and PFC (*blue*) for the monkey-side image features model (*left*), human-side image features model (*middle*) and the monkey-side joint features model (*right*). Asterisks represent statistical significance of the comparison (* is p < 0.05, ** is p <0.005, etc in a Wilcoxon rank sum test comparing model correlations of neurons of the corresponding brain regions). The results paint a dynamic picture of the dissociation between visual and motor regions in the natural task, which is presumably driven by the correlation between visual and motor features around various key events. See Figure S4.

**Figure S4.**
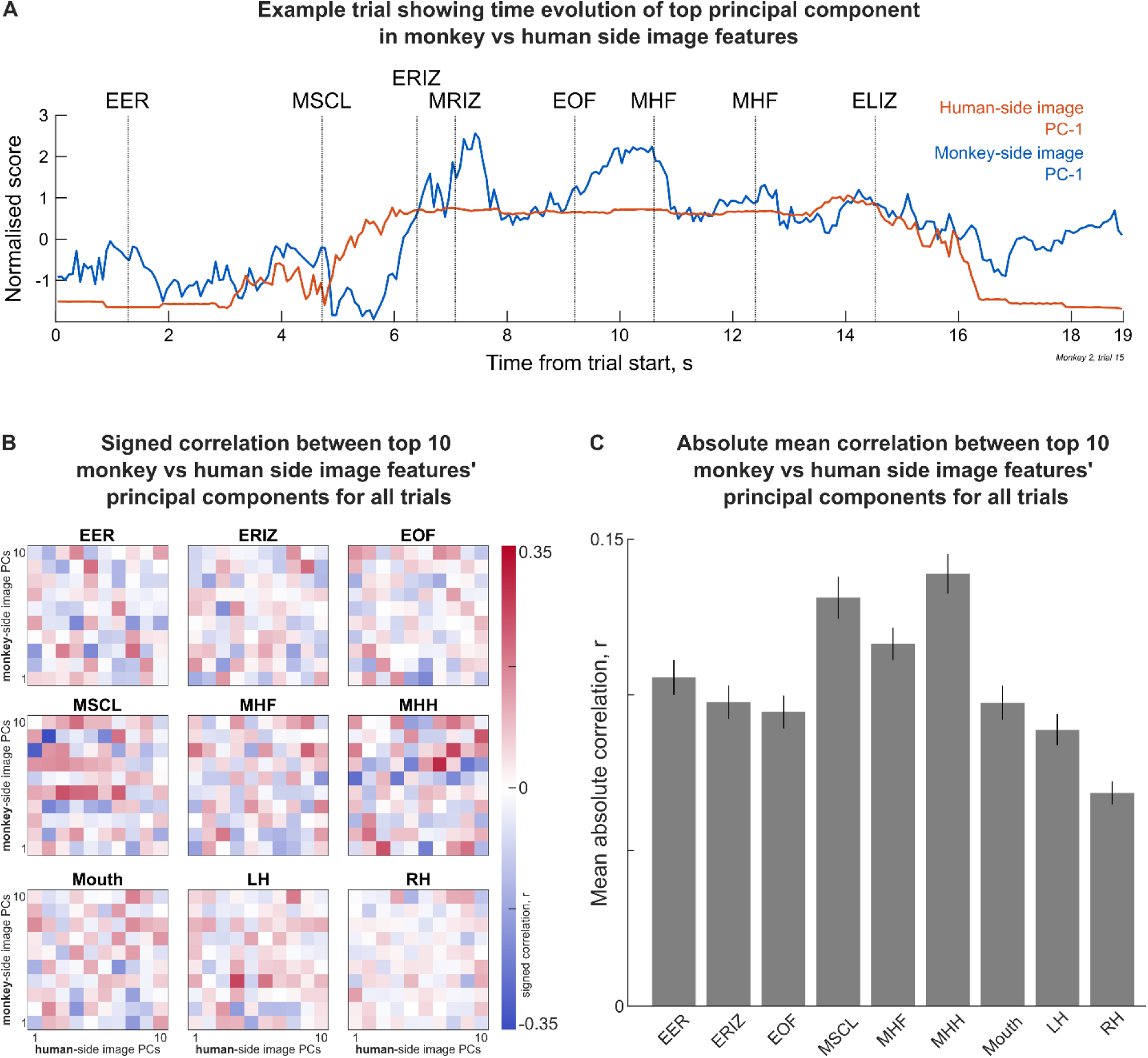
Visual and motor feature correlations in the natural task (A) Example naturalistic task trial in Monkey M2, showing the time course of the first principal component (PC) of the human-side image features (orange trace, explained variance = 47%) and first PC of the monkey-side image features (*blue trace*, explained variance = 22%). *Dotted vertical lines* indicate event times during the entire trial, as following: EER = Experimenter Entered Room, MSCL = Monkey Start Climb Log, ERIZ = Experimenter Reached Interaction Zone, MRIZ = Monkey Reached Interaction Zone, EOF = Experimenter Offered Food, MHF = Monkey Held Food, ELIZ = Experimenter Left Interaction Zone. (B) Colormap of pair-wise, signed correlation between the top-10 PCs of human-side visual features (along x-axis) and monkey-side visual features (along y-axis) in the natural task, evaluated in a window [-0.5 0.5] s relative to the event onset. (C) Bar plot of the average absolute correlation across all pairs of human-side and monkey-side PCs for each natural task event. It can be seen that some events have larger correlation between the human and monkey-side features than others: EOF (Experimenter Offered Food) has low correlation, presumably because the monkey was looking at the experimenter while eating the food. By contrast, MSCL (Monkey Start Climb Log) had a high correlation, presumably because the monkey and experimenter were moving towards each other at the same time.

